# GENOME REPORT: High-quality genome assemblies of 15 Drosophila species generated using Nanopore sequencing

**DOI:** 10.1101/267393

**Authors:** Danny E. Miller, Cynthia Staber, Julia Zeitlinger, R. Scott Hawley

## Abstract

The *Drosophila* genus is a unique group containing a wide range of species that occupy diverse ecosystems. In addition to the most widely studied species, *Drosophila melanogaster*, many other members in this genus also possess a well-developed set of genetic tools. Indeed, high-quality genomes exist for several species within the genus, facilitating studies of the function and evolution of cis-regulatory regions and proteins by allowing comparisons across at least 50 million years of evolution. Yet, the available genomes still fail to capture much of the substantial genetic diversity within the *Drosophila* genus. We have therefore tested protocols to rapidly and inexpensively sequence and assemble the genome from any Drosophila species using single-molecule sequencing technology from Oxford Nanopore. Here, we use this technology to present high-quality genome assemblies of 15 Drosophila species: 10 of the 12 originally sequenced Drosophila species (*ananassae, erecta, mojavensis, persimilis, pseudoobscura, sechellia, simulans, virilis, willistoni*, and *yakuba*), four additional species that had previously reported assemblies (*biarmipes, bipectinata, eugracilis*, and *mauritiana*), and one novel assembly (*triauraria*). Genomes were generated from an average of 29x depth-of-coverage data that after assembly resulted in an average contig N50 of 4.4 Mb. Subsequent alignment of contigs from the published reference genomes demonstrates that our assemblies could be used to close over 60% of the gaps present in the currently published reference genomes. Importantly, the materials and reagents cost for each genome was approximately $1,000 (USD). This study demonstrates the power and cost-effectiveness of long-read sequencing for genome assembly in Drosophila and provides a framework for the affordable sequencing and assembly of additional Drosophila genomes.

## INTRODUCTION

The early availability of high-quality genome assemblies for 12 species of Drosophila fostered many studies in evolution and comparative genomics, reinforcing Drosophila’s role as a primary model organism (Adams *et al*. 2000; Drosophila 12 Genomes Consortium 2007). Since the publication of these sequences, improvements have been made to the original 12 genomes (Hoskins *et al*. 2015) and the genomes for several additional species have been reported (c.f. Ometto *et al*. 2013; c.f. Nolte *et al*. 2013; Chiu *et al*. 2013; Allen *et al*. 2017). Although the quality and value of these genomes is high, the cost and effort required to assemble new genomes remains prohibitive for many laboratories. These issues, as well as the difficulty of assembling repetitive or low-complexity regions using short-read technology alone, must be overcome before we can rapidly increase the number of sequenced species.

Long-read, or third-generation, sequencing technology promises to simplify genome assembly by generating individual reads longer than many of the repetitive or low-complexity regions that have complicated genome assembly in the past (Chaisson *et al*. 2015). Indeed, long-read data for the *Drosophila melanogaster* reference genome stock ISO-1 generated on a Pacific Biosciences RSII were released in 2014 (Kim *et al*. 2014). These data were used to assemble a high-quality *D. melanogaster* genome with a contig N50 of 21 Mb (contig N50 is a measure of genome continuity in which half of the genome is contained in overlapping DNA segments, or contigs, larger than the value given), demonstrating that long reads could be used to generate highly contiguous genome assemblies in Drosophila (Berlin *et al*. 2015).

Genome assemblies using Oxford Nanopore sequencing technology have been reported for several species, including *Saccharomyces cerevisiae* (Salazar *et al*. 2017), *Arabidopsis thaliana* (Michael *et al*. 2017), *Caenorhabditis elegans* (Tyson *et al*. 2017), and *Homo sapiens* (Jain *et al*. 2018). This technology measures changes in current as a molecule (currently either DNA or RNA) passes through a small pore in a membrane. It has several advantages over other long-read technologies, including low DNA input requirements, ease of library preparation, and no theoretical limit to the length of a sequencing read. Although there are concerns about a high error rate with this technology, there are methods to mitigate this concern (Walker *et al*. 2014; Simpson *et al*. 2017; Vaser *et al*. 2017). For example, polishing using the original long-read data or data from another technology such as Illumina can correct SNP and indel errors by aligning reads to the assembled genome and identifying sites where modifications to the assembly result in resolution of a SNP or indel (Walker *et al*. 2014; Simpson *et al*. 2017). The cost of Nanopore sequencing is also attractive for small to medium-sized genome assembly—as of early 2018, up to 15 Gb of data could be generated on a single flow cell under ideal conditions for approximately $1,000 USD. Because of the advantages and the relatively low cost, we wondered if we could create a high-quality, non-scaffolded genome assembly from reads generated using a single flow cell.

Here, we report the sequencing and assembly of 15 *non-melanogaster* Drosophila species: *ananassae, biarmipes, bipectinata, erecta, eugracilis, mauritiana, mojavensis, persimilis, pseudoobscura, sechellia, simulans, triauraria, virilis, willistoni*, and *yakuba* (Figure 1). Nanopore sequencing and assembly of *D. melanogaster*, including genome scaffolding using additional technologies, is presented in a cosubmitted manuscript (Solares *et al*., this issue). These 15 species were sequenced to an average depth of coverage of 29x and an average read length of 5.9 kb. We rapidly assembled each genome using minimap2 and miniasm(Li 2016; 2018), resulting in an average contig N50 of 4.4 Mb. Assemblies were polished using either Nanopore data alone, Illumina short-read data alone, or a combination of both. Polishing with both Nanopore and Illumina data resulted in genomes containing on average 97.7% of all single-copy genes expected to be present in metazoans, consistent with current Drosophila reference assemblies. In addition, for the 10 species included here as part of our analysis of the original 12 genomes project, 97.8% of transcripts either fully or partially aligned to our assembled contigs, only 1.7% fewer transcripts than aligned to the current published reference genomes.

**Figure 1.**
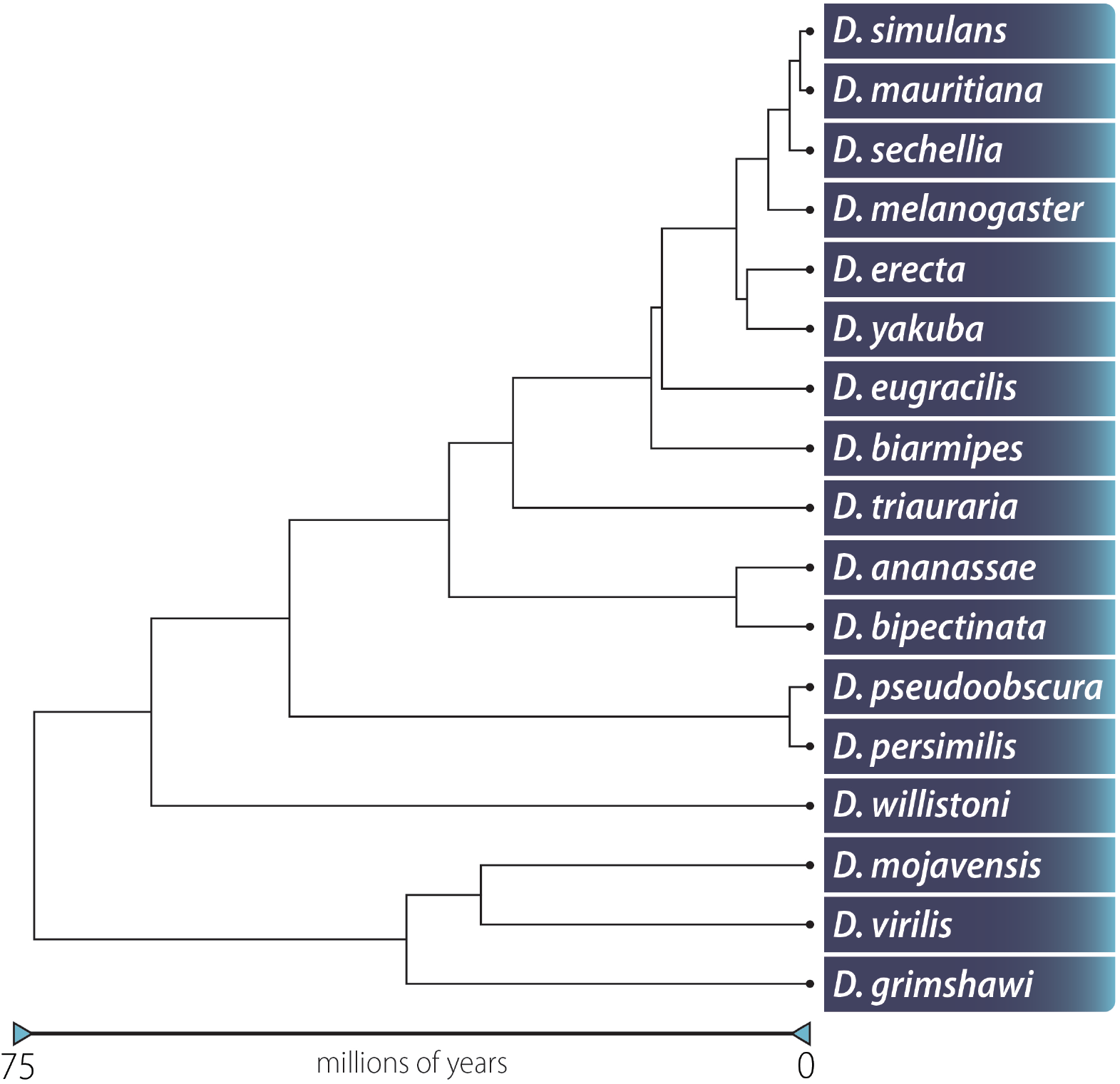
Phylogenetic tree of flies sequenced in this report including two species (*D. melanogaster* and *D. grimshawi*) not sequenced here but that were part of the original 12 genomes project (Drosophila 12 Genomes Consortium 2007). Adapted from Thomas and Hahn (2017).

Finally, after mapping contigs from currently published reference genomes to determine how many gaps could be closed using our highly contiguous assemblies, we estimate that an average of 61% of gaps in the currently published reference genomes could be resolved using our data. While the quality of each assembly was high, the overall materials and reagents cost was relatively low at approximately $1,000 USD per genome. This study demonstrates that it is feasible to generate a high-quality, low-cost genome for any member of the Drosophila genus and provides a framework by which additional Drosophila genomes may be assembled.

## MATERIALS and METHODS

### Stocks and reference genomes

All stocks used in this study (Figure 1, Table 1) are available at the National Drosophila Species Stock Center (http://blogs.cornell.edu/drosophila/). Selected stocks were used either in the original 12 genomes project or for subsequent published genome assemblies. Originally, eight stocks were obtained from an inhouse collection at the Stowers Institute (*ananassae, erecta, mojavensis, sechellia, simulans, virilis, pseudoobscura*, and *yakuba*) and eight stocks (*biarmipes*, *bipectinata*, *eugracilis*, *kikkawai*, *mauritiana*, *miranda*, *persimilis*, and *willistoni*) were obtained from the stock center when it was located at UCSD. Unfortunately, two of the eight stocks from the stock center (*kikkawai* and *miranda*) were found to be mislabeled. We were subsequently able to determine that the stock labeled *kikkawai* was in fact *triauraria*, but we were unable to determine what species the incorrectly labeled *miranda* stock was and thus removed it from our analysis. All flies were kept in standard cornmeal-agar bottles at 25 °C. The *D. mauritiana* reference genome was downloaded from http://www.popoolation.at/mauritiana_genome/index.html, and other reference genomes were downloaded from either FlyBase (ftp://ftp.flybase.net/genomes/) or GenBank (https://www.ncbi.nlm.nih.gov/assembly) (Table S1).

**Table 1.** Stocks sequenced in this study.

| Species | Stock #<sup>1</sup> |
| --- | --- |
| *D. ananassae* | 14024-0371.13 |
| *D. biarmipes* | 14023-0361.02 |
| *D. bipectinata* | 14024-0381.07 |
| *D. erecta* | 14021-0224.01 |
| *D. eugracilis* | 14026-0451.02 |
| *D. mauritiana* | 14021-0241.01 |
| *D. mojavensis* | 15081-1352.22 |
| *D. persimilis* | 14011-0111.01 |
| *D. pseudoobscura* | 14011-0121.94 |
| *D. sechellia* | 14021-0248.01 |
| *D. simulans* | 14021-0251.006 |
| *D. triauraria* | 14028-0691.9 |
| *D. virilis* | 15010-1051.87 |
| *D. willistoni* | 14030-0811.00 |
| *D. yakuba* | 14021-0261.01 |

<sup>1</sup> Stocks were obtained from the Drosophila Stock Center when it was located at the University of California San Diego. The stock center is now located at Cornell University.

### DNA isolation and quantification

Either virgin or non-virgin females were collected under CO2 and immediately frozen at -70 °C for at least 1 hr before DNA isolation. DNA was isolated using either the Qiagen Blood & Cell Culture DNA Mini Kit or by a phenol protocol (Table S1). For the Blood & Cell Culture kit (column-based isolation) 60-100 frozen females were placed in two 1.5-mL Eppendorf tubes and frozen in liquid nitrogen before being homogenized using a pestle in 250 μl of Buffer G2 with 200 μg/ml RNase A. 700 μl of Buffer G2 with RNase A, and 50 μl of 20 mg/mL proteinase K was then added to each tube, followed by incubation at 50°C for 2 hr. Each tube was spun at 5k RPM for 5 min, then the supernatant was removed and placed in a new 1.5-mL Lo-Bind Eppendorf tube and vortexed for 10 seconds. The supernatant from both tubes was then transferred onto the column and allowed to flow through via gravity. The column was washed 3x with wash buffer and eluted twice with 1 mL of elution buffer into 2 separate 1.5-mL Lo-Bind tubes. 700 μl of isopropanol was added and mixed via inversion before being spun at 14,000 RPM for 15 min at 4°C. The supernatant was removed and the pellet was washed with freshly prepared 70% ethanol, then centrifuged at 14,000 RPM for 10 min at 4°C. The supernatant was removed and 25 μl of nuclease-free water was added to each tube and allowed to sit at room temperature for 2 hr. Both tubes were then combined and stored at 4°C.

For phenol-based DNA extractions, 60-100 frozen flies were transferred to a 2-mL Kontes Dounce homogenizer (VWR #KT885300-0002) on ice in 1 mL homogenization buffer (0.1 M NaCL, 30 mM Tris-HCl pH 8.0, 10 mM EDTA, 0.5% Triton X-100). Flies were dounced 4-5x with each pestle (first the looser pestle A, then the tighter pestle B). Homogenate was transferred to a 1.5-mL Eppendorf tube on ice using a wide-bore pipet tip. Homogenizer was rinsed with 500 μL homogenization buffer and combined with homogenate. Debris was pelleted by centrifugation for 1 min at 500x g at 4°C.

A wide-bore pipet tip was used to transfer supernatant containing nuclei in suspension to a clean tube. Nuclei were pelleted by centrifugation for 5 min at 2000x g at 4°C. Supernatant was carefully decanted and the pelleted nuclei were resuspended in 200 μL homogenization buffer by pipetting with a large-bore tip. Nuclei were transferred to a clean tube and lysed by adding 1.268 mL extraction buffer (0.1 M Tris-HCl pH 8.0, 0.1 M NaCl, 20 mM EDTA), 1.5 μL proteinase K (20 mg/mL, Life Technologies AM2548), and 30 μL 10% SDS. Lysis was aided by gently swirling and rocking, followed by incubation at 37°C for 2-4 hr without agitation.

Lysed nuclei were extracted twice with an equal volume of phenol/chloroform/isoamyl alcohol pH 8.0 (Amresco K169-100ML). Extraction was mixed gently on a rotator for 5 min then centrifuged 5 min at 5000x g at room temperature. The upper aqueous phase was transferred to a clean tube and the extraction repeated as above with chloroform (Sigma C2432). The final upper aqueous phase was transferred to a clean tube and precipitated by adding 0.1 volume 3M NaOAc (mixed gently) and 2 volumes absolute ethanol (mixed by gentle inversion). A wide-bore pipette tip was used to transfer the DNA (a white, stringy clump) to a clean tube. (Note: If high-quality low-retention tips are not available, it is recommended to Sigmacote the tips to prevent the gDNA from adhering to the plastic.) Excess ethanol was removed by pipetting. DNA was washed with 500 μL 70% ethanol and centrifuged briefly at low speed with a tabletop centrifuge. Supernatant was removed by pipetting and DNA was allowed to air dry ~10 minutes before resuspending in 100 μL TE pH 8.0 at 4 °C overnight.

### Nanopore library preparation, sequencing, and base calling

Libraries were prepared using the Ligation Sequencing Kit 1D (Oxford Nanopore) either according to or with slight modifications to the manufacturer’s protocol. To begin the prep an average of 2.3 μg of DNA was used, which is higher than the 400 ng recommended by the manufacturer. Water or TE was added to DNA for a total volume of 46 μL. For 4 of 21 library preps (Table S1), the FFPE repair and dA-Tailing steps were combined in the following reaction mix: 46.5 μL of genomic DNA in TE, 3.5 μL of UltraII EP Buffer (NEB), 3.5 μL of FFPE DNA Repair Buffer (NEB), 3 μL of UltraII EP Enzyme (NEB), 3 μL of FFPE Repair Mix (NEB), and 0.5 μL of 100x NAD+ (NEB). The combined reaction was prepared in a 200-μL PCR tube and run at 20°C for 1 hr followed by 65°C for 30 min in a thermocycler. After cleanup and adapter ligation, 75 μL of library (note that all 15 μl of adapter-ligated DNA, not the 12 μL recommended by the manufacturer, was included in the final library), including Library Loading Beads, were loaded onto an R9.4 flow cell containing at least 800 active pores and run for 48-72 hr, or until no pores were available, on a MinION sequencer (Oxford Nanopore). Flow cells were restarted 3-6 times during a run in order to increase the number of pores in strand at any given time. Separate flow cells were used for each species. Nine species each utilized a single flow cell while two flow cells were used for the following six species: *D. virilis*, because of its large genome size; *D. simulans*, because of low read output on the first run; *D. bipectinata, D. erecta*, *D. eugracilis*, and *D. mojavensis*, because of a substandard library kit on the first run that produced fewer and shorter reads than expected (Table S1). Base calling was completed using Albacore Sequencing Pipeline Software version 2.1.0 (Oxford Nanopore) with default settings, and fastq files for either all reads or only those that passed the default quality filter (quality score ≥7) were combined for assembly and polishing (Table 2, Table S2).

**Table 2.** Base-called reads used for genome assembly.^*1*^

| Species | Depth of coverage |  |  | All reads |  | Reads >1 kb |  | Reads >10 kb |  | Longest read (bp) |
| --- | --- | --- | --- | --- | --- | --- | --- | --- | --- | --- |
|  | All reads | Number of reads >1 kb | Number of reads >10 kb | Total number of reads | Average length (bp) | Number of reads | Average length (bp) | Number of reads | Average length (bp) |  |
| *D. ananassae* | 44.8 | 42.9 | 20.1 | 2,085,829 | 4,227 | 1,409,289 | 5,990 | 252,071 | 15,690 | 110,391 |
| *D. biarmipes* | 28.8 | 27.5 | 12.9 | 1,375,651 | 4,102 | 861,668 | 6,232 | 166,579 | 15,155 | 101,492 |
| *D. bipectinata* | 22.6 | 20.5 | 9.9 | 1,590,181 | 2,909 | 716,348 | 5,861 | 109,688 | 18,473 | 279,705 |
| *D. erecta* | 31.7 | 30.6 | 17.5 | 1,022,546 | 4,924 | 658,736 | 7,373 | 151,078 | 18,358 | 176,712 |
| *D. eugracilis* | 22.5 | 21.2 | 8.2 | 1,424,426 | 3,617 | 905,481 | 5,369 | 117,316 | 15,923 | 152,351 |
| *D. mauritiana* | 32.6 | 32.1 | 17.1 | 852,775 | 6,045 | 717,219 | 7,066 | 175,669 | 15,402 | 93,106 |
| *D. mojavensis* | 45.4 | 44.0 | 23.5 | 1,471,959 | 5,129 | 1,063,822 | 6,871 | 189,302 | 20,659 | 245,241 |
| *D. persimilis* | 34.5 | 33.6 | 15.5 | 1,386,759 | 4,910 | 1,072,290 | 6,179 | 199,330 | 15,304 | 113,218 |
| *D. pseudoobscura* | 33.3 | 31.7 | 13.8 | 1,417,469 | 3,936 | 905,571 | 5,868 | 154,234 | 15,000 | 104,401 |
| *D. sechellia* | 23.9 | 23.8 | 16.9 | 390,359 | 10,200 | 366,174 | 10,828 | 110,941 | 25,347 | 254,031 |
| *D. simulans* | 30.2 | 30.0 | 25.1 | 389,278 | 12,393 | 311,889 | 15,346 | 140,196 | 28,532 | 309,608 |
| *D. triauraria* | 18.8 | 18.7 | 13.1 | 409,756 | 9,634 | 379,831 | 10,338 | 120,668 | 22,876 | 238,837 |
| *D. virilis* | 22.2 | 21.8 | 11.9 | 1,209,939 | 5,969 | 1,017,016 | 6,976 | 189,572 | 20,402 | 282,795 |
| *D. willistoni* | 28.5 | 26.2 | 12.9 | 1,868,768 | 3,132 | 923,706 | 5,816 | 158,505 | 16,757 | 100,073 |
| *D. yakuba* | 21.7 | 21.5 | 12.6 | 509,806 | 7,277 | 462,732 | 7,947 | 103,820 | 20,699 | 195,439 |
| **Average** | **29.4** | **28.4** | **15.4** | **1,131,520** | **5,894** | **771,222** | **6,931** | **154,466** | **18,693** | n/a |

<sup>1</sup>All reads listed had quality scores $\geq 7$ .

**Table 3.** Assembly statistics for published assemblies using scaffold values and contig statistics of minimap2 assemblies after polishing with Racon three times followed by Pilon three times.

|  |  | Published assembly |  |  |  |  |  | Minimap2 assemblies using only “pass” reads |  |  |  |  |
| --- | --- | --- | --- | --- | --- | --- | --- | --- | --- | --- | --- | --- |
| Species | Estimated genome size (Mb) | Version | Assembly size (Mb) | Number of scaffolds | Scaffold N50 (Mb) | Number of scaffolds >100 kb | Number of scaffolds >1 Mb | Assembly size (Mb) | Number of contigs | Contig N50 (Mb) | Number of contigs >100 kb | Number of contigs >1 Mb |
| *D. ananassae* | 196.6 | 1.05 | 231.0 | 13,749 | 4.6 | 100 | 29 | 188.6 | 371 | 2.6 | 147 | 44 |
| *D. biarmipes* | 195.6 | 2.0 | 169.4 | 5,523 | 3.4 | 95 | 32 | 184.6 | 666 | 2.8 | 224 | 32 |
| *D. bipectinata* | 204.6 | 2.0 | 167.3 | 5,500 | 0.7 | 124 | 35 | 160.5 | 570 | 0.6 | 354 | 26 |
| *D. erecta* | 158.9 | 1.05 | 152.7 | 5,124 | 18.7 | 38 | 8 | 127.7 | 58 | 16.6 | 29 | 21 |
| *D. eugracilis* | 228.9 | 2.0 | 156.9 | 4,946 | 1.0 | 150 | 38 | 156.4 | 547 | 1.0 | 256 | 42 |
| *D. mauritiana* | 157.9 | 1.0 | 117.7 | 16 | 21.1 | 12 | 7 | 131.9 | 272 | 4.7 | 64 | 28 |
| *D. mojavensis* | 166.3 | 1.04 | 193.8 | 6,841 | 24.8 | 51 | 12 | 162.1 | 122 | 5.0 | 86 | 39 |
| *D. persimilis* | 197.1 | 1.3 | 188.4 | 12,838 | 1.9 | 142 | 30 | 165.7 | 415 | 3.5 | 146 | 35 |
| *D. pseudoobscura* | 167.7 | 3.04 | 152.7 | 4,790 | 12.5 | 25 | 13 | 160.5 | 361 | 3.0 | 143 | 33 |
| *D. sechellia* | 166.7 | 1.3 | 166.6 | 14,730 | 2.1 | 101 | 23 | 133.4 | 109 | 7.4 | 59 | 23 |
| *D. simulans* | 159.6 | 2.02 | 125.0 | 7,619 | 23.5 | 8 | 6 | 132.2 | 76 | 7.7 | 61 | 24 |
| *D. triauraria* | 210.2 | n/a | n/a | n/a | n/a | n/a | n/a | 170.5 | 482 | 0.72 | 339 | 34 |
| *D. virilis* | 325.4 | 1.06 | 206.0 | 13,530 | 10.2 | 79 | 22 | 165.9 | 141 | 4.1 | 83 | 40 |
| *D. willistoni* | 205.4 | 1.05 | 235.5 | 14,838 | 4.5 | 80 | 38 | 197.7 | 490 | 1.5 | 270 | 49 |
| *D. yakuba* | 170.7 | 1.05 | 165.7 | 8,122 | 21.8 | 60 | 8 | 141.1 | 111 | 5.2 | 68 | 32 |

### Illumina library preparation and sequencing

DNA for Illumina sequencing was prepared from 10 males using a Qiagen Blood and Tissue Kit (Table S3). Briefly, flies were frozen at -70 °C for at least 1 hr before DNA extraction following the manufacturer’s instructions. Genomic DNA was sonicated using the Covaris S220 and libraries were constructed on a Perkin Elmer Sciclone G3 NGS Workstation using the KAPA HTP Library Prep kit (KAPA Biosystems, Cat. No. KK8234) and NEXTflex DNA barcodes (Bioo Scientific, Cat No. NOVA-514104). Post-amplification size selection was performed on all libraries using a Pippin Prep (Sage Science). Resulting libraries were quantified using an Agilent 2100 Bioanalyzer plus an Invitrogen Qubit 2.0 Fluorometer and then pooled. Sequencing was performed on an Illumina NextSeq 500 instrument as 150 bp on a high-output, paired-end flow cell. Illumina NextSeq Real Time Analysis version 2.4.11 and bcl2fastq2 v2.18 were run to demultiplex sequencing reads and to generate FASTQ files.

### Genome assembly, polishing, and quality evaluation

Minimap2 (Version 2.1-r311) (Li 2018) and miniasm (version 0.2-r168-dirty) (Li 2016) were used to assemble either all reads or only those reads with quality scores ≥7. Polishing was completed using either Racon (Vaser *et al*. 2017) or Pilon (Walker *et al*. 2014) alone or in combination (Table 4, Tables S5-S8). Dot plots were generated with nucmer and mummerplot (Delcher *et al*. 1999); for clarity, only contigs or scaffolds >100 kb were plotted. Assemblies using only reads that passed quality filter were subjected to either four iterations of polishing with Racon, six iterations of polishing with Pilon, or three iterations of polishing with Racon followed by three iterations of polishing with Pilon. BUSCO version 2.0.1 (Simão *et al*. 2015) was used to evaluate assembly quality for all genomes using the metazoan_odb9 database, which contains 978 single-copy genes likely to be present in any metazoan genome (Table 4, Tables S5-S8). For the 10 genomes assembled in this report that were part of the original Drosophila 12 genomes, transcripts were downloaded from FlyBase and aligned to each assembly and each reference genome using BLAST (Altschul *et al*. 1997). For each species and both genome types (assembly and reference), those transcripts for which at least 90% of the transcript aligned to the genome with at least 95% identity were counted as proper alignments (Table S9). To perform SNP calling, Illumina reads were aligned using bwa version 0.7.17-r1188 (Li and Durbin 2009) to assembled genomes that had been polished with Racon and Pilon. Samtools was then used to identify SNPs and indels. Only those SNPs and indels with quality scores >220 were counted (Table 5).

**Table 4.** BUSCO scores^*1*^ reveal assembly quality.

| Species | Published assembly | minimap2 | minimap2 →<br>Racon x1 | minimap2 →<br>Racon x4 | minimap2 →<br>Pilon x1 | minimap2 →<br>Pilon x6 | minimap2 →<br>Racon x3 →<br>Pilon x3 |
| --- | --- | --- | --- | --- | --- | --- | --- |
| *D. ananassae* | 98.2 | 1.3 | 88.3 | 91.6 | 74.7 | 91.8 | 98.2 |
| *D. biarmipes* | 98.6 | 3.8 | 88.3 | 92 | 77.1 | 94.7 | 98.7 |
| *D. bipectinata* | 98.2 | 1.7 | 74.9 | 80.1 | 67.2 | 87.9 | 93.9 |
| *D. erecta* | 98.6 | 2.5 | 89.9 | 90.4 | 79.5 | 95.5 | 98.6 |
| *D. eugracilis* | 98.5 | 0.8 | 82.6 | 86.7 | 70.7 | 92.7 | 97.9 |
| *D. mauritiana* | 98.6 | 1.4 | 91 | 94.6 | 75.9 | 95 | 98.7 |
| *D. mojavensis* | 98.2 | 0.5 | 82.5 | 88.7 | 66.9 | 93.5 | 98 |
| *D. persimilis* | 96.6 | 0.4 | 80.3 | 85.3 | 62.1 | 90.9 | 98 |
| *D. pseudoobscura* | 97.0 | 1.7 | 84.1 | 88.8 | 72.3 | 93.9 | 98 |
| *D. sechellia* | 97.2 | 1.5 | 91.3 | 92.1 | 75.3 | 95.2 | 98.7 |
| *D. simulans* | 98.6 | 2.7 | 91.3 | 95.6 | 77.5 | 94.4 | 98.6 |
| *D. triauraria* | NA | 3.1 | 82.9 | 85.0 | 73.5 | 87.9 | 93.8 |
| *D. virilis* | 97.5 | 1.1 | 84.9 | 89.3 | 70.9 | 93.6 | 97.7 |
| *D. willistoni* | 98.4 | 0.6 | 79.7 | 82.8 | 72.2 | 92.1 | 98.1 |
| *D. yakuba* | 98.5 | 1.3 | 86.7 | 91.2 | 73.7 | 95.5 | 98.4 |

<sup>1</sup>Only complete BUSCO scores for the minimap2 assembly using reads that passed filter ( $\geq 7$ ) are shown. All values are percentages. Higher scores suggest better assembly.

**Table 5.**
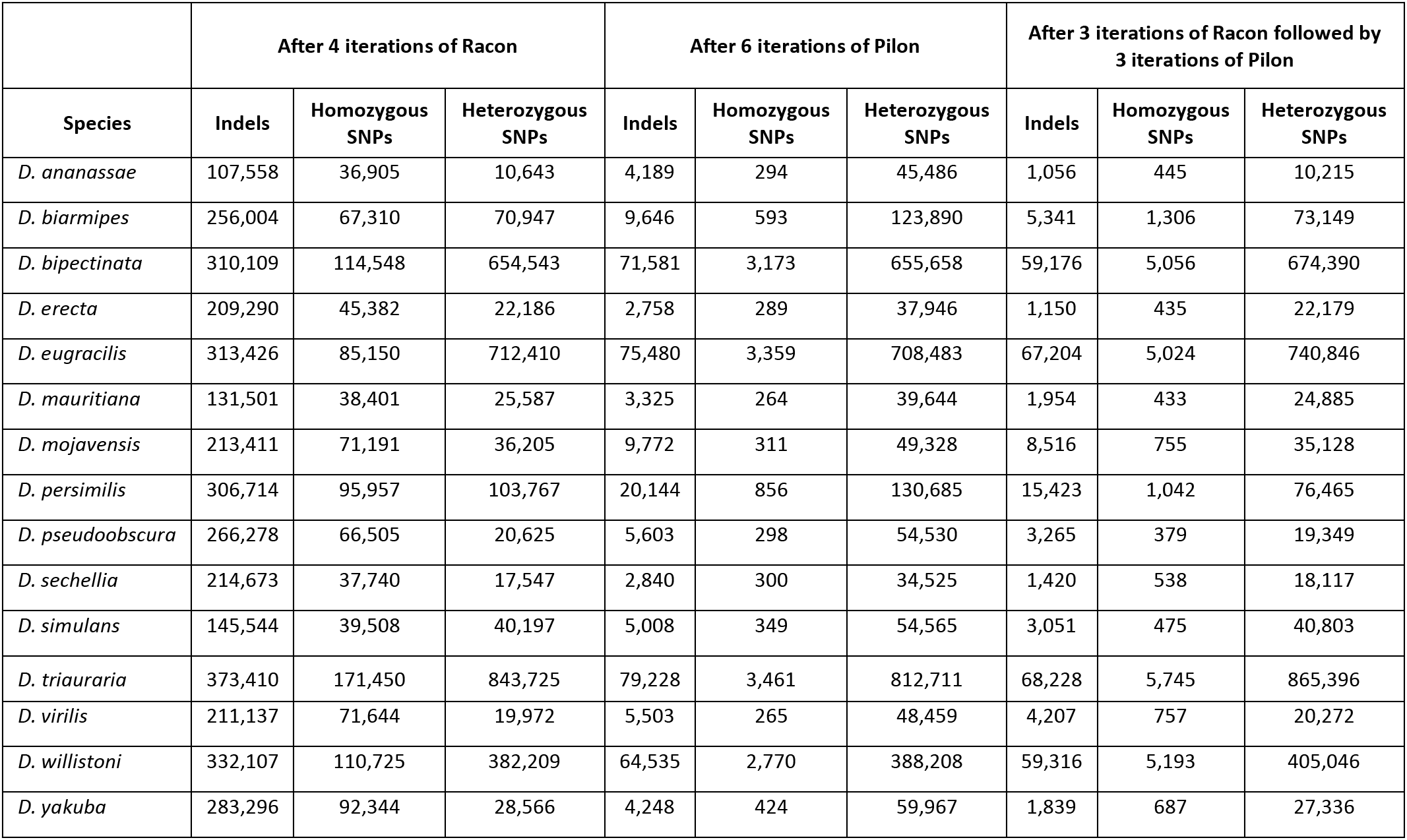
Number of single nucleotide and indel polymorphisms after polishing.

| Species | After 4 iterations of Racon |  |  | After 6 iterations of Pilon |  |  | After 3 iterations of Racon followed by 3 iterations of Pilon |  |  |
| --- | --- | --- | --- | --- | --- | --- | --- | --- | --- |
|  | Indels | Homozygous SNPs | Heterozygous SNPs | Indels | Homozygous SNPs | Heterozygous SNPs | Indels | Homozygous SNPs | Heterozygous SNPs |
| *D. ananassae* | 107,558 | 36,905 | 10,643 | 4,189 | 294 | 45,486 | 1,056 | 445 | 10,215 |
| *D. biarmipes* | 256,004 | 67,310 | 70,947 | 9,646 | 593 | 123,890 | 5,341 | 1,306 | 73,149 |
| *D. bipectinata* | 310,109 | 114,548 | 654,543 | 71,581 | 3,173 | 655,658 | 59,176 | 5,056 | 674,390 |
| *D. erecta* | 209,290 | 45,382 | 22,186 | 2,758 | 289 | 37,946 | 1,150 | 435 | 22,179 |
| *D. eugracilis* | 313,426 | 85,150 | 712,410 | 75,480 | 3,359 | 708,483 | 67,204 | 5,024 | 740,846 |
| *D. mauritiana* | 131,501 | 38,401 | 25,587 | 3,325 | 264 | 39,644 | 1,954 | 433 | 24,885 |
| *D. mojavensis* | 213,411 | 71,191 | 36,205 | 9,772 | 311 | 49,328 | 8,516 | 755 | 35,128 |
| *D. persimilis* | 306,714 | 95,957 | 103,767 | 20,144 | 856 | 130,685 | 15,423 | 1,042 | 76,465 |
| *D. pseudoobscura* | 266,278 | 66,505 | 20,625 | 5,603 | 298 | 54,530 | 3,265 | 379 | 19,349 |
| *D. sechellia* | 214,673 | 37,740 | 17,547 | 2,840 | 300 | 34,525 | 1,420 | 538 | 18,117 |
| *D. simulans* | 145,544 | 39,508 | 40,197 | 5,008 | 349 | 54,565 | 3,051 | 475 | 40,803 |
| *D. triauraria* | 373,410 | 171,450 | 843,725 | 79,228 | 3,461 | 812,711 | 68,228 | 5,745 | 865,396 |
| *D. virilis* | 211,137 | 71,644 | 19,972 | 5,503 | 265 | 48,459 | 4,207 | 757 | 20,272 |
| *D. willistoni* | 332,107 | 110,725 | 382,209 | 64,535 | 2,770 | 388,208 | 59,316 | 5,193 | 405,046 |
| *D. yakuba* | 283,296 | 92,344 | 28,566 | 4,248 | 424 | 59,967 | 1,839 | 687 | 27,336 |

### Alignment of reference contigs to Nanopore assemblies

Each reference genome was broken into contigs by separating scaffolds at every position that contained an N, and BLAST (Altschul *et al*. 1997) was used to align these to contigs from our Nanopore assembly using a custom script. We first aligned only reference contigs larger than 10 kb from reference scaffolds that contained no N’s (that had no gaps) and counted how many of these reference contigs mapped with an identity >99% to at least one assembled contig. We then aligned only reference contigs that came from reference scaffolds with two or more contigs (meaning the reference scaffold had at least one gap) one reference scaffold at a time. A gap was assumed to be filled if at least two reference contigs mapped to the same assembled contig.

### Data availability

Illumina data generated for this project are available at the National Center for Biotechnology Information (https://www.ncbi.nlm.nih.gov/) under project PRJNA427774, Nanopore reads that passed the default Albacore filter are available under project PRJNA471302. Illumina and Nanopore data for *D. triauraria* are available under project PRJNA473618. Scripts used in this project and genomes assembled in this project can be found on GitHub at https://github.com/danrdanny/Drosophila15GenomesProject/. Original data underlying this manuscript can be accessed from the Stowers Original Data Repository at http://www.stowers.org/research/publications/libpb-1269.

## RESULTS AND DISCUSSION

### Sequencing

Nanopore sequencing runs on a flow cell containing a maximum of 2,048 pores through which singlestranded DNA passes. These 2,048 pores are contained within 512 channels, with four pores per channel. Currently, only one pore in a channel—512 total pores—are available for sequencing at a time. The sequencing control software, called MinKNOW, consigns each pore on a flow cell into one of four groups (often called mux groups) based on its performance level. Each mux group is run for a predetermined amount of time (8 hours by default), at which point sequencing switches to the next group of pores because pore quality deteriorates over time.

For this study, we generally ran a sequencing reaction for 24 hours, allowing each of the pores in groups 1-3 to run for 8 hours (pores in group 4 are not used during a run). We then stopped and restarted the flow cell, allowing pores to be reorganized into new groups, which also allows pores originally assigned to the unused group 4 to be moved back into groups 1-3. We then monitored the flow cell and restarted again as the number of active pores decreased during a run, once again reorganizing pores into new groups. Because the number of active pores deteriorates over time and the amount of data output by a flow cell drops dramatically after 24 hours, there is an advantage to keeping as many pores as possible actively sequencing, or “in strand,” at a time.

Preliminary testing in our lab revealed that using the manufacturer-recommended 400 ng of input DNA for a 1D library prep resulted in fewer than 50% of active pores in strand. This low number of active pores translated into low total data output by the flow cell. We found that starting a 1D library prep with 1–10 μg of DNA resulted in substantially more pores in strand during a sequencing run and gave higher data yields. We therefore performed all library preps in this study using 1-10 μg of starting material. Sequencing 15 Drosophila species in this manner yielded a total of 23 million sequence reads (Table S2).

After base calling, these 23 million reads, with an average read length of 4,302 bp, yielded ~99 billion bases sequenced at an average depth of coverage of 35x (Table S2). The base calling software, Albacore, separates reads into ‘pass’ and ‘fail’ bins by default, where any read with a quality score ≥7 is identified as a read that passed filter. The 76% of reads and 85% of total bases that passed filter had an average read length of 5,894 bp and an average depth of coverage of 29x, while those that did not pass filter had an average read length of 2,680 bp (Table 2, Table S2).

To determine whether variations in certain steps of the protocol might provide increased data output, we tested three different methods of DNA extraction and preparation. First, we compared column-based and phenol extraction (see Materials and Methods). While column-based methods are more convenient and safe, phenol extractions may reduce DNA shearing and loss. Second, in an attempt to reduce DNA shearing even further through reduced pipetting, we performed phenol extractions followed by a shortened library preparation protocol in which the FFPE repair and dA-tailing steps were combined. For reads passing filter, the average read length of samples prepared by a column-based method was 4,013 bp. Average read length sharply increased to 8,931 bp in phenol extractions that followed library prep instructions, while phenol-extracted samples that combined the FFPE repair and dA-tailing steps showed an average read length of 10,389 bp. These results suggest that combining these two library preparation steps is potentially useful, but the greatest increase in read length resulted from careful phenol-based DNA isolation.

### Genome assembly and statistics

De novo assembly of 15 genomes can be computationally expensive since the software typically used to assemble long error-prone sequencing reads (for example Canu (Koren *et al*. 2017), or hybrid assemblers, such as DBG2OLC (Ye *et al*. 2016), which uses both short high-quality reads and long error-prone reads) requires a large number of CPU hours to complete. This may be cost-prohibitive for some labs. Alternatively, mapping and assembly using minimap2 (an alignment program that can map long sequences with a high error rate to a reference database) followed by miniasm (a *de novo* assembler for long error-prone reads) has been shown to be highly efficient and less time-intensive. Indeed, minimap itself showed a fivefold improvement in mapping over other commonly used mapping software (Li 2016; 2017).

A study comparing assembly quality of the *D. melanogaster* ISO-1 reference genome using different combinations of these assembly approaches found that assemblies using minimap alone gave larger contigs (N50 3.9 Mb) than Canu alone (N50 3.0 Mb) but smaller contigs than DBG2OLC alone (N50 9.9 Mb) or a merged Canu and DBG2OLC assembly (N50 18.6 Mb) (Solares *et al*., co-submitted). This suggests that assembly statistics from minimap2 and miniasm may be comparable to some more computationally intensive long-read assembly software (Table 4). We therefore chose to perform assembly using only minimap2 and miniasm. Each of the 15 assemblies was completed in under 1 hour using 32 CPUs on a computer with 512 Gb of RAM available. While assembly statistics are similar between these assemblies and a Canu assembly, it is important to note that the quality of these assemblies is much lower because Canu contains an assembly error correcting step (discussed below). The addition of an equivalent error correction step to a minimap2 assembly does slightly increase assembly time, but remains less than that of Canu alone.

For each species, sequenced reads were combined into two fastq files, one containing all reads and one containing only those reads that passed filter. Separate genome assemblies were then completed for each category of reads and compared. Assemblies using only reads that passed filter had larger contig N50 values in 9 of 15 cases and fewer overall contigs in 10 of 15 cases than assemblies completed using all sequencing reads (Table 3, Table S4). Because of the moderately improved N50 values and lower contig numbers, we decided to proceed with our analysis using only assemblies generated by higher quality reads. Assembly quality varied greatly among the 15 genomes, with an average contig N50 of 4.4 Mb, a maximum of 16.6 Mb (*D. erecta*), and a minimum of 0.6 Mb (*D. bipectinata*). For each species, assembly resulted in genome sizes that were smaller than the expected genome size (Bosco *et al*. 2007; Gregory and Johnston 2008; Hjelmen and Johnston 2017), with repetitive sequence likely accounting for the lower values than expected. We also compared assembly statistics for each of our *de novo* assemblies to the published assembly of each species and found that in all cases our contig N50 values were higher than those of the published assemblies but lower than the scaffold N50 values (scaffold N50 is a measure similar to contig N50, in which half of the genome is contained in linked DNA segments, or scaffolds, larger than the value given) in all but three cases (Table 3). Published assemblies contain both contigs (contiguous DNA segments) and scaffolds (one or more contigs generally separated by gaps where the DNA sequence is unclear), while the assemblies presented here contain only contigs.

We next generated dot plots to better understand how our assemblies compare to the reference genome for the 10 species sequenced as part of the original Drosophila 12 genomes project (Drosophila 12 Genomes Consortium 2007). Although overall our assemblies correlated well with the published reference assemblies, we did observe a few inversions, rearrangements, or duplications in each plot (Figure S1). For example, a small inversion is seen between the *D. yakuba* reference chromosome *2L* and our assembly contig utg000002l, similar to other large inversions that have been reported to be segregating in this population (Llopart *et al*. 2002). Indeed, large rearrangements and inversions are observed in several of the assemblies. It is possible that these changes may have occurred in the stock after being initially isolated for genome sequencing, they may be due to assembly errors in the initial assemblies as they were generated using relatively low-coverage data, or the stocks we sequenced, while labeled as such, may not have been the exact stock used for sequencing by the 12 genomes consortium. We also observe differences in repetitive, poorly assembled regions of the genome. For example, the X-axis of the *D. sechellia* and *D. persimilis* plots are longer than the Y-axis, likely because our assemblies have collapsed repetitive regions that the published reference genomes have not.

### Assembly polishing and evaluation

While the N50 statistic has utility as a measure of genome contiguity, it is not necessarily a good indicator of genome quality. A highly contiguous genome may in fact contain many errors that make it difficult to use for downstream analysis such as gene finding or SNP calling. We did not expect our Nanopore-only assemblies to be of high quality because raw Nanopore reads have an observed error rate of 10-15% (Li 2016; Simpson *et al*. 2017; Salazar *et al*. 2017). It is, however, possible to improve assembly quality through one or more rounds of polishing, which is a process that improves assembly quality using the original sequencing reads, reads from another technology with a relatively low error rate (e.g., Illumina data), or a combination of both. We therefore sought to evaluate both the quality of our initial minimap2 assemblies as well as our assemblies using various combinations of polishing techniques.

Assembly quality was evaluated using BUSCO, which searches assemblies for highly conserved genes generally present in a single copy in any genome (Simão *et al*. 2015). For example, BUSCO analysis of published reference genomes for the 15 species sequenced in this study revealed that at least 96.6% of 978 highly conserved genes were present in at least one copy in each genome (Table 4, Table S4). We ran BUSCO on all 15 of our minimap2 assemblies with no polishing and found an average of 1.6% (min: 0.4%, max: 3.8%) of genes were present in at least one copy, suggesting that our initial assemblies, although highly contiguous, were indeed of poor quality. This is likely due to the high error rate of the reads used to generate the assemblies, resulting in frame shifts leading to premature stop codons within open reading frames.

We then performed multiple iterations of polishing using either Racon or Pilon (Walker *et al*. 2014; Vaser *et al*. 2017). Racon, which uses only base-called Nanopore reads for polishing, improved average BUSCO scores from 1.6% to 85.2% after a single iteration (min: 74.9%, max: 91.3%). Repeat iterations of Racon alone improved scores only slightly, reaching an average of 88.9% after four iterations (min: 80.1%, max: 95.6%) (Figure 2A, Table 4, Table S5). Pilon alone, which uses only Illumina data for polishing, improved average BUSCO scores from 1.6% to 72.6% after a single iteration (min: 62.1%, max: 79.5%), and to 93.0% after six iterations (min: 87.9%, max: 95.5%) (Figure 2B, Table 4, Table 6). While BUSCO scores tended to increase with more iterations, it is important to note that BUSCO scores fell for 7 of 15 assemblies between the 3^rd^ and 4^th^ iterations of Racon, suggesting that repeat iterations of polishing may have negative impacts on assembly quality. Alignment of Illumina data to Racon-polished genomes demonstrated that polishing using Racon alone failed to completely correct poly-N indels. Furthermore, while Pilon was run for 6 consecutive iterations, there was not significant improvement in scores after 3 iterations, demonstrating a limit to polishing using a single software application.

**Figure 2.**
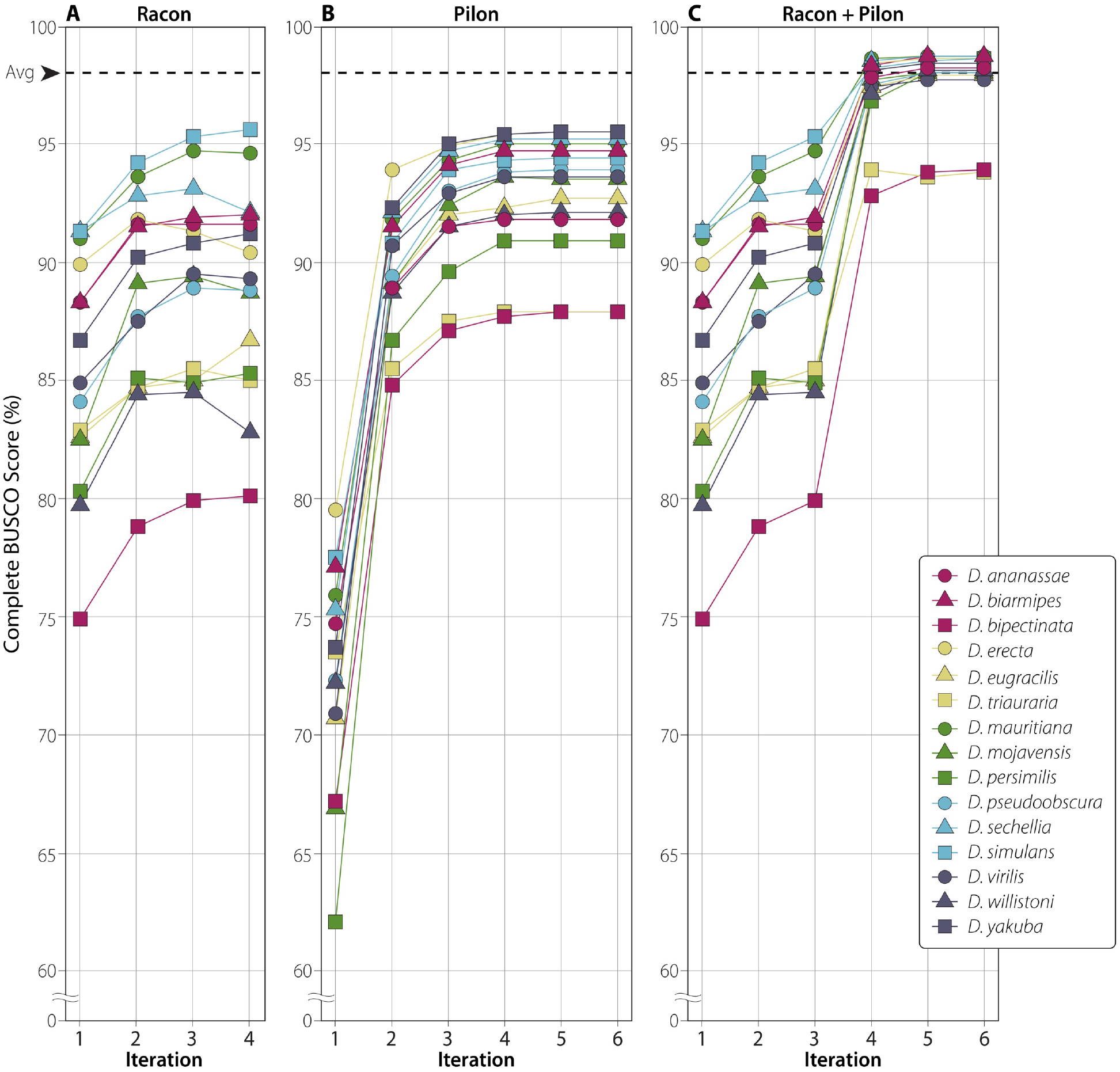
Polishing improves assembly quality. Average BUSCO score for these 15 assembled genomes was 1.6% before polishing. The dotted line in all panels represents the average BUSCO score for all 14 published reference genomes (Table 4). Full polishing results can be found in Tables S5-S8. **(A)** Complete BUSCO scores for four iterations of Racon alone. **(B)** Complete BUSCO scores for six iterations of Pilon alone. **(C)** Complete BUSCO scores shown for three iterations of Racon followed by three iterations of Pilon.

**Table 6.**
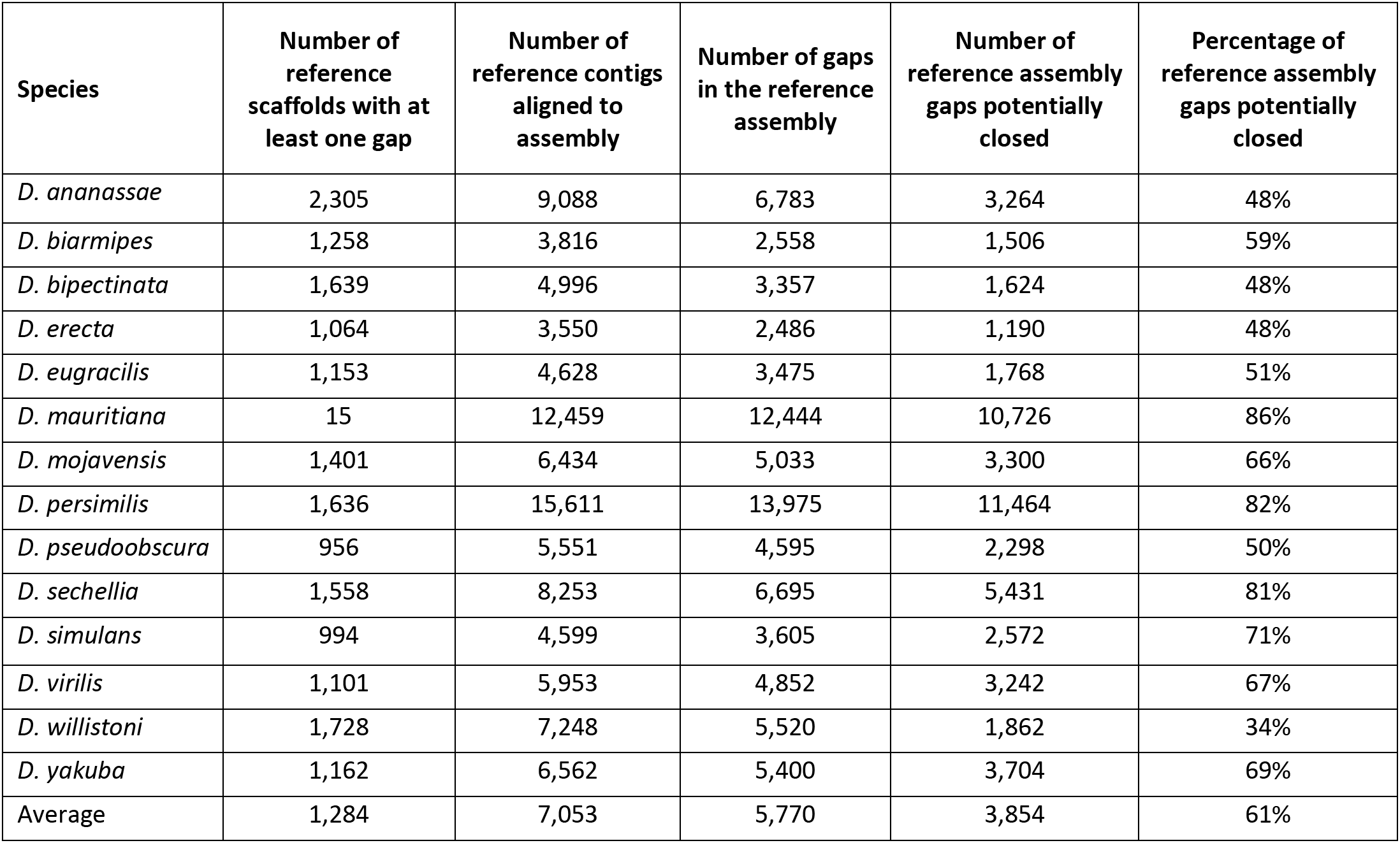
Number of singleton reference contigs that could be placed on the contigs assembled in this study and the number of gaps closed between reference contigs on scaffolds with one or more gaps.

Because both polishing techniques alone failed to achieve BUSCO scores equal to or better than the published reference genomes, we then polished using a combination of both Racon and Pilon. We first attempted to run Pilon and Racon in combination, one after the other (e.g., Racon, Pilon, Racon, Pilon, etc.), but found that while BUSCO scores improved with each iteration of Pilon, they then fell with each iteration of Racon (Table S7). Because BUSCO scores plateau around 3 iterations of Racon or Pilon (see above), we wondered if combining the two approaches in tandem would result in improved assembly quality. We therefore ran three iterations of Racon followed by three iterations of Pilon and found a sustained improvement in BUSCO scores, reaching an average of 97.7% (min: 93.8%, max: 98.7%) (Figure 2C, Table 4, Table S8). Importantly, this combination of polishing resulted in BUSCO scores consistent with the currently published reference genomes for 13 of the previously sequenced species. Finally, to determine if currently available gene models properly mapped to our assemblies, we mapped transcripts for the 10 assemblies that are available on FlyBase to both our assembled genomes and the currently published reference genomes. We found that on average 97.8% of all transcripts mapped to our assembled genomes, while 99.6% mapped to the published reference genome (Table S9). This suggests that while the BUSCO scores of our genomes suggest similarity between our assemblies and the reference, a small number of genes that are not considered highly conserved by BUSCO may have been improperly assembled using our approach.

To determine if high levels of heterozygosity of the stocks used for sequencing explained the lower BUSCO scores observed in the *D. bipectinata* and *D. triauraria* stocks (average of 93.9%), we aligned Illumina data from each stock to the assembly generated from three iterations of Racon and Pilon and called SNPs and indels using SAMtools (Li *et al*. 2009). On average, we observed 63,702 indels, and 769,893 heterozygous SNPs in these two stocks, compared to an average of 13,365 indels and 116,445 heterozygous SNPs in the remaining 13 stocks. This high number of SNP and indel polymorphisms suggests a high level of heterozygosity in the stocks with low BUSCO scores. However, we also observed a high number of SNP and indels in both the *D. eugracilis* and *D. willistoni* stocks, two assemblies with BUSCO scores near 98%, suggesting that heterozygosity for SNPs and indels alone does not fully explain the lower quality scores observed in *D. bipectinata* and *D. triauraria*.

These observations led us to ask why a large number of heterozygous SNPs were observed in four of our lines. Because no effort was made to isogenize these stocks before sequencing, it is possible that the high number of SNPs and indels in these lines is simply a consequence of selection for heterozygosity in these lines. However, this hypothesis would predict high levels of heterozygosity in the other sequenced stocks, which is not observed. Another possibility is that that these four lines were crossed (either purposely or by contamination) with another stock in order to increase the fitness of the stock, which would have introduced the large amount of heterozygosity we observe.

Finally, we wondered if polishing would merge or separate contigs and thus affect assembly statistics. We found that polishing with each program independently did, on average, increase the total assembly size and contig N50 of our 15 assemblies (Table S9). Furthermore, the total number of contigs decreased for 6 of 15 species and increased for none after polishing with Racon, while total contig number remained unchanged after polishing with Pilon. Overall, this suggests that polishing using Racon or Pilon had minimal impact on assembly statistics, but clearly improved assembly quality as determined by BUSCO scores.

### Long-read data can close gaps in reference genomes

Other long-read sequencing technologies, such as PacBio, have previously been used to improve existing reference genomes (Jiao *et al*. 2017), and visual inspection of dot plots used to compare our assemblies with the reference genome suggest that our contigs could be used to link large reference scaffolds together (Figure S1). To test if our highly contiguous genomes could be used to fill gaps in the reference assemblies, we broke each reference genome into individual contigs and aligned those contigs to our assembled genomes. We performed two types of alignment. First, we identified reference contigs larger than 10 kb that contained no N’s, or unknown sequence (these contigs are generally labeled as scaffolds in many reference genomes, which may cause confusion) and identified how many of those could be placed on our assemblies with >99% identity. In doing so, we found that each reference genome contained an average of 205 singleton contigs larger than 10 kb, and that on average 95% of these could be placed on a larger contig in our assemblies (Table 6). (*D. mauritiana* was excluded from this analysis because its genome assembly is based on release 5 of the *D. melanogaster* genome and thus contains only one contig larger than 10 kb with no gaps.)

We then identified scaffolds from each reference genome that contained two or more contigs separated by N’s and found that the 14 reference genomes contain an average of 1,284 scaffolds with two or more contigs (min: 15, max: 2,305). For each reference scaffold, individual reference contigs were mapped to the assembled genome and the gap between mapped reference contigs was determined. A single gap was considered closed when reference contigs from the same reference scaffold were mapped to a single assembled contig. This mapping revealed that, on average, our assembled contigs closed 3,854 of 5,770, or 61%, of gaps in each of the reference genomes (Table 6).

A specific example of how long, contiguous assemblies may assist with reference genome correction is scaffold_4845 in the *D. erecta* reference assembly. This 22.6-Mb reference scaffold contains 75 contigs and 112,933 N’s. We broke this scaffold into its 75 contigs (numbered 0-74) and were able to place 39 of these reference contigs (numbered 23-61) onto contig utg000010l, a single 17.4-Mb contig from our assembly. These 39 contigs were placed in the same order as they appear in the reference assembly (Figure 3), but with important exceptions. Specifically, three reference contigs were placed within larger reference contigs. For example, contig 53 is a 79,102-bp contig that was placed in the middle of contig 52, a 339,371-bp contig. Similarly, contig 39 was placed within 40, and contig 60 was placed within 61. All six contigs have >99% identity to the positions that they were aligned to, suggesting either that the reference assembly incorrectly identified these contigs as unique segments of the genome or that our assembly collapsed nearby regions of the genome with high identity into single segments.

**Figure 3.**
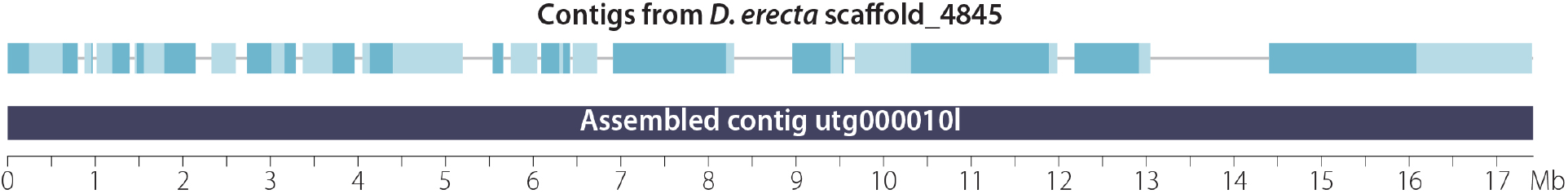
Gaps in the reference genome assembly can be closed using long-read data. This example shows that a 17.4-Mb contig (utg000010l) from our assembly (bottom) closes the gaps (top, gray lines) among 38 contigs (top, shaded boxes) from the *D. erecta* reference scaffold (scaffold_4845), potentially resolving 3.7 Mb of sequence.

### Sequencing costs and conclusions

A major goal of this project was to determine if it was possible to create low-cost, high-quality genome assemblies of *non-melanogaster* Drosophila species using Nanopore sequencing. We are able to estimate an expected cost for each of our Drosophila genomes using publicly available materials and reagent prices from early 2018. Here, we used approximately one flow cell per species; when purchased individually, a single flow cell is $900 (USD) and when purchased as a pack of 48, a single flow cell is $500 (USD). A 1D library kit is $599 (USD) and will make six libraries, at a cost of $100 per genome. Reagents include FFPE enzyme, dA-tailing enzyme, ligase, and magnetic beads for cleanup. To simplify, we assume $50 in reagent and other costs per genome. Therefore, for Nanopore sequencing alone, the costs for materials and reagents range from $650-$1,050 depending on the flow cell cost. We also generated Illumina 150-bp paired-end data for polishing at an average of 64x depth of coverage (Table S3). While short read sequencing costs vary widely, we assume a cost of $250 per Drosophila genome is a reasonable estimate. Therefore, we estimate that overall sequencing and assembly costs for a high-quality Drosophila reference genome to range from $900-$1,300 (USD).

In summary, we have generated high-quality genome assemblies for 15 species of Drosophila at a materials and reagents cost of approximately $1000 (USD). Relatively low-coverage Nanopore data resulted in genome assemblies with an average contig N50 value of 4.4 Mb, and polishing with Racon and Pilon resulted in average BUSCO scores of 97.7%, a score comparable to currently published reference genomes (Figure 2, Table 3, Table 4). That we were able to generate 15 different genome assemblies at relatively low cost and in a short period of time suggests a new approach researchers may take when studying other species within the *Drosophila* genus and suggests that genome assembly using a long-read technology should now be considered the standard for studies of new Drosophila species.

It is now also feasible to consider what effort, if any, should be put forth in attempts to assemble and analyze a substantially larger number of genomes from across the entire *Drosophila* genus. The *Drosophila* genus represents at least 50 million years of evolution (Tamura *et al*. 2004; Obbard *et al*. 2012; O’Grady and DeSalle 2018) and spans a broad range of ecosystems, with the unique advantage of having a highly developed genetic toolkit available for many of its members (Ashburner *et al*. 2005; Gratz *et al*. 2013; Perkins *et al*. 2015; Stern *et al*. 2017). Low-cost, high-quality genome assemblies of hundreds of well-described species would foster studies of genome conservation and evolution orders of magnitude more detailed than any before and on a scale not possible in other similarly related species. Such work would provide a solid foundation to ensure the next 100 years of Drosophila research are as fruitful as the first 100 years have been.

## ACKNOWLEDGEMENTS

We thank A. Bernardo Carvalho for identifying that the *D. kikkawai* stock originally included in this report was incorrectly labeled, Amir Yassin for identifying the likely source of the mix-up and for providing information on species identification in the *montium* species group, and Patrick O’Grady for confirming that the stock is indeed *D. triauraria*. We also thank Justin Blumenstiel, Rob Unckless and members of the Bachtrog laboratory and Hawley laboratory for helpful comments, discussions, and suggestions, as well as Angela Miller for assistance with editing and figure preparation. DEM has received support in the form of free flow cells from Oxford Nanopore as part of the Oxford Nanopore Educational Initiative. RSH is an American Cancer Society Research Professor.

---

**Figure S1.**
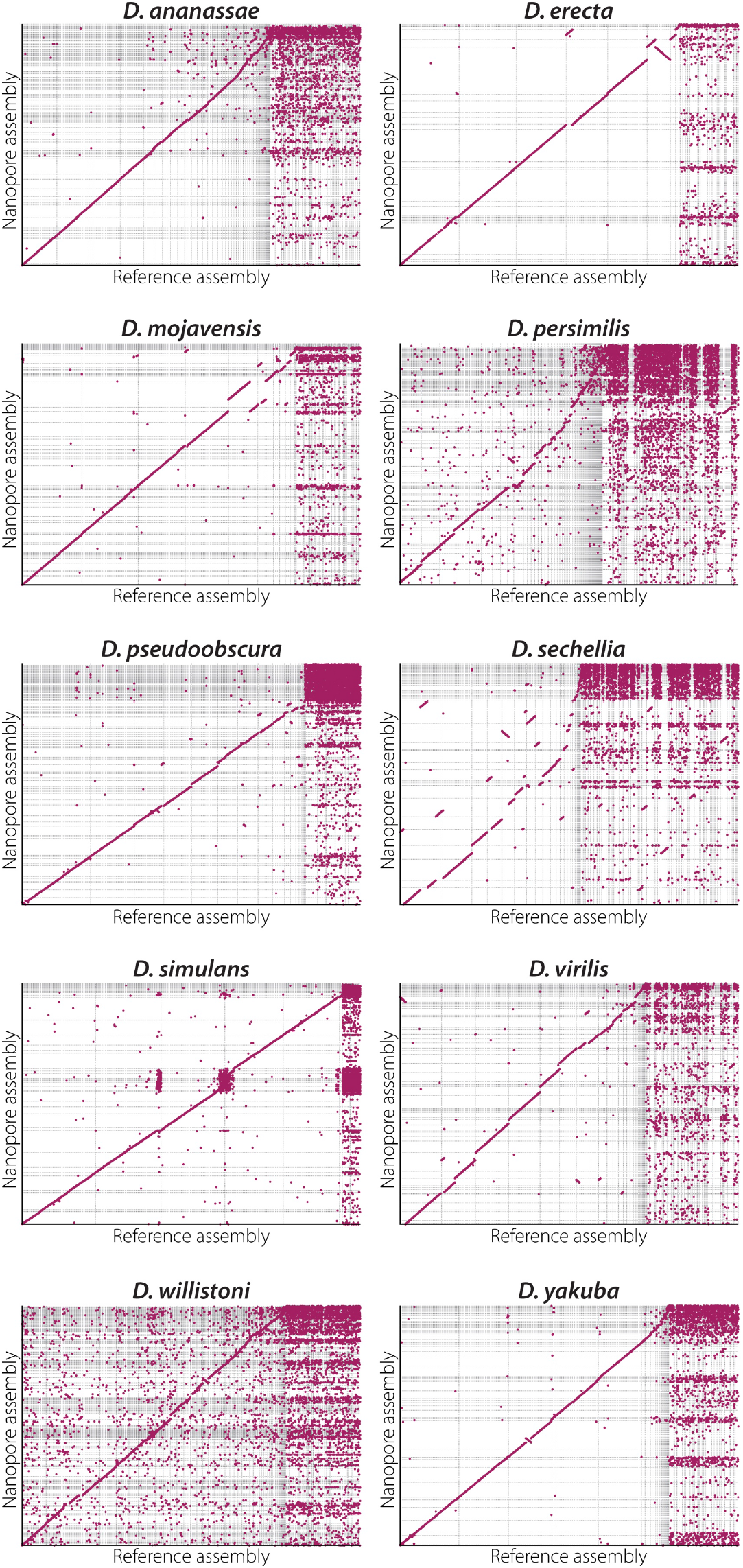
Dot plots comparing Nanopore assemblies presented in this project (y-axis) with reference genome assemblies (x-axis) for 10 species sequenced as part of the 12 genomes project (Drosophila 12 Genomes Consortium 2007). A single dot or line is plotted for a sequence from one genome that is similar to sequence in the other genome. Dots or lines with a positive slope indicate the sequences are oriented in the same direction, while a negatively sloping line represents sequences oriented in opposite directions. Gridlines define scaffolds in the reference genome or contigs from the Nanopore assemblies.

**Table S1:** Stocks sequenced in this study.

**Table S2:** Detail of all sequencing reads.

**Table S3:** Summary of Illumina NextSeq data used for polishing.

**Table S4:** Assembly statistics and BUSCO scores for reference genomes and unpolished genomes assembled using either all reads or only those that passed filter.

**Table S5:** Assembly statistics and BUSCO scores for all genomes after polishing with Racon up to 4 times.

**Table S6:** Assembly statistics and BUSCO scores for all genomes after polishing with Pilon up to 6 times.

**Table S7:** Assembly statistics and BUSCO scores for all genomes after polishing with Racon 3 times followed by polishing with Pilon 3 times.

**Table S8:** Assembly statistics and BUSCO scores for all genomes after iteratively polishing with Racon followed by Pilon.

**Table S9:** Transcripts from published reference genomes mapped to Nanopore assemblies.

**Table S10:** Differences in assembly statistics between initial assembly and either Racon-only or Pilon-only polished assemblies.

## LITERATURE CITED

Adams, M. D., S. E. Celniker, R. A. Holt, C. A. Evans, J. D. Gocayne et al., 2000 The genome sequence of Drosophila melanogaster. Science 287: 2185–2195.

Allen, S. L., E. K. Delaney, A. Kopp, and S. F. Chenoweth, 2017 Single-Molecule Sequencing of the *Drosophila serrata* Genome. G3 (Bethesda) 7: 781–788.

Altschul, S. F., T. L. Madden, A. A. Schäffer, J. Zhang, Z. Zhang et al., 1997 Gapped BLAST and PSI-BLAST: a new generation of protein database search programs. Nucleic Acids Research 25: 3389–3402.

Ashburner, M., K. Golic, and R. S. Hawley, 2005 Drosophila: A Laboratory Handbook. Cold Spring Harbor Laboratory Press, Cold Spring Harbor, NY.

Berlin, K., S. Koren, C.-S. Chin, J. P. Drake, J. M. Landolin et al., 2015 Assembling large genomes with single-molecule sequencing and locality-sensitive hashing. Nat Biotechnol 33: 623–630.

Bosco, G., P. Campbell, J. T. Leiva-Neto, and T. A. Markow, 2007 Analysis of Drosophila Species Genome Size and Satellite DNA Content Reveals Significant Differences Among Strains as Well as Between Species. Genetics 177: 1277–1290.

Chaisson, M. J. P., R. K. Wilson, and E. E. Eichler, 2015 Genetic variation and the de novo assembly of human genomes. Nature Publishing Group 16: 627–640.

Chiu, J. C., X. Jiang, L. Zhao, C. A. Hamm, J. M. Cridland et al., 2013 Genome of *Drosophila suzukii*, the spotted wing drosophila. G3 (Bethesda) 3: 2257–2271.

Delcher, A. L., S. Kasif, R. D. Fleischmann, J. Peterson, O. White et al., 1999 Alignment of whole genomes. Nucleic Acids Research 27: 2369–2376.

Drosophila 12 Genomes Consortium, 2007 Evolution of genes and genomes on the Drosophila phylogeny. Nature 450: 203–218.

Gratz, S. J., A. M. Cummings, J. N. Nguyen, D. C. Hamm, L. K. Donohue et al., 2013 Genome Engineering of Drosophila with the CRISPR RNA-Guided Cas9 Nuclease. Genetics 194: 1029–1035.

Gregory, T. R., and J. S. Johnston, 2008 Genome size diversity in the family Drosophilidae. Heredity 101: 228–238.

Hjelmen, C. E., and J. S. Johnston, 2017 The mode and tempo of genome size evolution in the subgenus Sophophora. (I. V. Sharakhov, Ed.). PLoS ONE 12: e0173505.

Hoskins, R. A., J. W. Carlson, K. H. Wan, S. Park, I. Mendez et al., 2015 The Release 6 reference sequence of the *Drosophila melanogaster* genome. Genome Research 25: 445–458.

Jain, M., S. Koren, K. H. Miga, J. Quick, A. C. Rand et al., 2018 Nanopore sequencing and assembly of a human genome with ultra-long reads. Nat Biotechnol 1–16.

Jiao, Y., P. Peluso, J. Shi, T. Liang, M. C. Stitzer et al., 2017 Improved maize reference genome with single-molecule technologies. Nature Publishing Group 546: 524–527.

Kim, K. E., P. Peluso, P. Babayan, P. J. Yeadon, C. Yu et al., 2014 Long-read, whole-genome shotgun sequence data for five model organisms. Sci. Data 1: 140045–10.

Koren, S., B. P. Walenz, K. Berlin, J. R. Miller, N. H. Bergman et al., 2017 Canu: scalable and accurate long-read assembly via adaptive k-mer weighting and repeat separation. Genome Research 27: 722–736.

Li, H., 2016 Minimap and miniasm: fast mapping and de novo assembly for noisy long sequences. Bioinformatics 32: 2103–2110.

Li, H., 2018 Minimap2: pairwise alignment for nucleotide sequences. Bioinformatics 3: 321.

Li, H., 2017 Minimap2: versatile pairwise alignment for nucleotide sequences. arXiv 1–5.

Li, H., and R. Durbin, 2009 Fast and accurate short read alignment with Burrows-Wheeler transform. Bioinformatics 25: 1754–1760.

Li, H., B. Handsaker, A. Wysoker, T. Fennell, J. Ruan et al., 2009 The Sequence Alignment/Map format and SAMtools. Bioinformatics 25: 2078–2079.

Llopart, A., S. Elwyn, D. Lachaise, and J. A. Coyne, 2002 Genetics of a difference in pigmentation between *Drosophila yakuba* and *Drosophila santomea*. Evolution 56: 2262–2277.

Michael, T. P., F. Jupe, F. Bemm, S. T. Motley, J. P. Sandoval et al., 2017 High contiguity *Arabidopsis thaliana* genome assembly with a single nanopore flow cell. bioRxiv 149997.

Nolte, V., R. V. Pandey, R. Kofler, and C. Schlötterer, 2013 Genome-wide patterns of natural variation reveal strong selective sweeps and ongoing genomic conflict in *Drosophila mauritiana*. Genome Research 23: 99–110.

Obbard, D. J., J. Maclennan, K.-W. Kim, A. Rambaut, P. M. O’Grady et al., 2012 Estimating Divergence Dates and Substitution Rates in the Drosophila Phylogeny. Mol. Biol. Evol. 29: 3459–3473.

Ometto, L., A. Cestaro, S. Ramasamy, A. Grassi, S. Revadi et al., 2013 Linking genomics and ecology to investigate the complex evolution of an invasive Drosophila pest. Genome Biology and Evolution 5: 745–757.

O’Grady, P. M., and R. DeSalle, 2018 Phylogeny of the Genus Drosophila. Genetics 209: 1–25.

Perkins, L. A., L. Holderbaum, R. Tao, Y. Hu, R. Sopko et al., 2015 The Transgenic RNAi Project at Harvard Medical School: Resources and Validation. Genetics 201: 843–852.

Salazar, A. N., A. R. Gorter de Vries, M. van den Broek, M. Wijsman, P. de la Torre Cortés et al., 2017 Nanopore sequencing enables near-complete *de novo* assembly of *Saccharomyces cerevisiae* reference strain CEN.PK113-7D. FEMS Yeast Res. 17:.

Simão, F. A., R. M. Waterhouse, P. Ioannidis, E. V. Kriventseva, and E. M. Zdobnov, 2015 BUSCO: assessing genome assembly and annotation completeness with single-copy orthologs. Bioinformatics 31: 3210–3212.

Simpson, J. T., R. E. Workman, P. C. Zuzarte, M. David, L. J. Dursi et al., 2017 Detecting DNA cytosine methylation using nanopore sequencing. Nature Methods 14: 1–7.

Stern, D. L., J. Crocker, Y. Ding, N. Frankel, G. Kappes et al., 2017 Genetic and Transgenic Reagents for *Drosophila simulans*, D. mauritiana, D. yakuba, D. santomea, and D. virilis. G3 (Bethesda) 7: 1339–1347.

Tamura, K., S. Subramanian, and S. Kumar, 2004 Temporal patterns of fruit fly (Drosophila) evolution revealed by mutation clocks. Mol. Biol. Evol. 21: 36–44.

Thomas, G., and M. Hahn, 2017 Drosophila 25 species phylogeny. Figshare.

Tyson, J. R., N. J. O’Neil, M. Jain, H. E. Olsen, P. Hieter et al., 2017 MinION-based long-read sequencing and assembly extends the *Caenorhabditis elegans* reference genome. Genome Research gr.221184.117.

Vaser, R., I. Sović, N. Nagarajan, and M. Šikić, 2017 Fast and accurate de novo genome assembly from long uncorrected reads. Genome Research 27: 737–746.

Walker, B. J., T. Abeel, T. Shea, M. Priest, A. Abouelliel et al., 2014 Pilon: an integrated tool for comprehensive microbial variant detection and genome assembly improvement. (J. Wang, Ed.). PLoS ONE 9: e112963.

Ye, C., C. M. Hill, S. Wu, J. Ruan, and Z. S. Ma, 2016 DBG2OLC: Efficient Assembly of Large Genomes Using Long Erroneous Reads of the Third Generation Sequencing Technologies. Sci. Rep. 6: 1–9.

